# A mast cell receptor is associated with fibrosis and itch in systemic sclerosis

**DOI:** 10.64898/2026.10.02.756290

**Authors:** Ruchita Kothari, Daniel H. Capuzzi, Collin B. Kilgore, Hyun Jong Oh, Saloni Patel, Anjana Srikumar, Gwendolyn Leatherman, Ami A. Shah, Jun Kang, Xinzhong Dong

## Abstract

Systemic sclerosis (SSc) is an autoimmune connective tissue disease characterized by symmetric fibrosis of the skin and multiple internal organs. The mechanisms that trigger early fibrotic changes remain unclear. Mast cells are known to infiltrate human SSc lesional skin in early disease stages, yet the role of mast cells in initiating fibrosis and pruritus in SSc remains debated, in part due to unclear pathways of mast cell activation. Here, we show that mast cells, via the receptor Mrgprb2, contribute to fibrosis in the bleomycin (BLM)-induced skin fibrosis mouse model. Mast cells accumulate in the skin of BLM-treated wild-type (WT) but not Mrgprb2-knockout *(Mrgprb2*^−/−^) mice. *Mrgprb2*^−/−^ mice show attenuated disease severity, with reduced collagen deposition and lower expression of matrix metalloproteinases and inflammatory markers associated with fibrosis. BLM treatment also increases pruritus in WT mice in the early stages of the model; an effect reduced in *Mrgprb2*^−/−^ mice. This is the first report of itch in this model of skin fibrosis, further linking it to clinical SSc, where pruritus is an early and prevalent symptom affecting nearly half of patients.In a parallel human cohort, lesional SSc skin harbors significantly more mast cells than paired non-lesional skin from the same patients, demonstrating that mast cell accumulation is specific to affected tissue. Lesional mast cell counts also track with skin disease severity, correlating with modified Rodnan skin score and patient-reported skin tightness. Together, these findings implicate Mrgprb2-mediated mast cell activity in both fibrosis and pruritus in SSc, highlighting mast cells as a candidate therapeutic target in this understudied disease process.

## Introduction

Systemic sclerosis, or SSc, is an autoimmune connective tissue disease characterized by progressive, multiorgan fibrosis. Cutaneous manifestations are among the most frequent diagnostic criteria and can alone warrant diagnosis of SSc^1^. While the mechanisms that initiate SSc remain unclear, early disease pathogenesis involves crosstalk between endothelial and immune cells. Endothelial cell dysfunction may recruit inflammatory infiltrate into affected blood vessels, while immune activation in turn exacerbates endothelial injury^2^. This sustained vascular damage drives differentiation and activation of myofibroblasts, which deposit excess collagen in the extracellular matrix^3^, driving the hallmark fibrosis of SSc. These immunologic mechanisms also likely interfere with the skin’s sensory compartment, manifesting clinically as pruritus. In SSc, pruritus emerges as an early symptom^4,5^, affecting 45-62% of patients^6^, and correlating with more extensive skin disease and worsening pulmonary, gastrointestinal, and mental health outcomes^7,8,9^. Despite its prevalence and clinical impact, the pathogenesis of pruritus in SSc remains markedly understudied, highlighted by the lack of any itch-focused studies in any preclinical mouse model of SSc.

Mast cells are tissue-resident innate immune cells best known for IgE-mediated allergic responses. Beyond this canonical pathway, mast cells also signal through Mrgprb2 and its human ortholog MRGPRX2, G protein-coupled receptors activated by cationic, amphipathic ligands such as neuropeptides and host defense peptides^10–12^. Mrgprb2/X2 signaling has been linked to pruritus and chronic inflammatory skin conditions, including atopic dermatitis, psoriasis, and urticaria^13^. Mast cells more broadly promote fibroblast proliferation and collagen deposition^14^, yet the role of Mrgprb2/X2 in this fibrotic process remains unclear. Mast cells are known to accumulate in skin biopsies of SSc patients^15^, and mast cell-mediated itch has been proposed to contribute to SSc pruritus^16^, though the extent of this contribution remains uncertain^17^. Given the capacity of Mrgprb2-expressing mast cells to drive both acute inflammation^18^ and chronic fibrosis, we hypothesized that Mrgprb2 may contribute to dermal fibrosis and itch in SSc. Here, we show that mast cells, via Mrgprb2, contribute to fibrosis and itch in mouse SSc, and that mast cells are associated with lesional skin and skin severity in human SSc.

## Results

### Mrgprb2 contributes to collagen deposition and skin fibrosis in a BLM model of SSc

To induce skin fibrosis, the bleomycin model of skin fibrosis was carried out by injecting WT and *Mrgprb2*^−/−^ mice with bleomycin (BLM) subcutaneously in the back skin^19^ over 21 days (**Figure 1a**). BLM-treated WT mice show increased skin thickness demonstrated by hematoxylin and eosin (H&E) stain (**Figure 1b**). To quantify collagen deposition, skin sections were stained with Masson’s trichrome (**Figure 1c**). Consistent with the H&E stain, collagen thickness is significantly increased in BLM-treated WT skin compared to *Mrgprb2*^−/−^ skin (**Figure 1d**).

**Figure 1.**
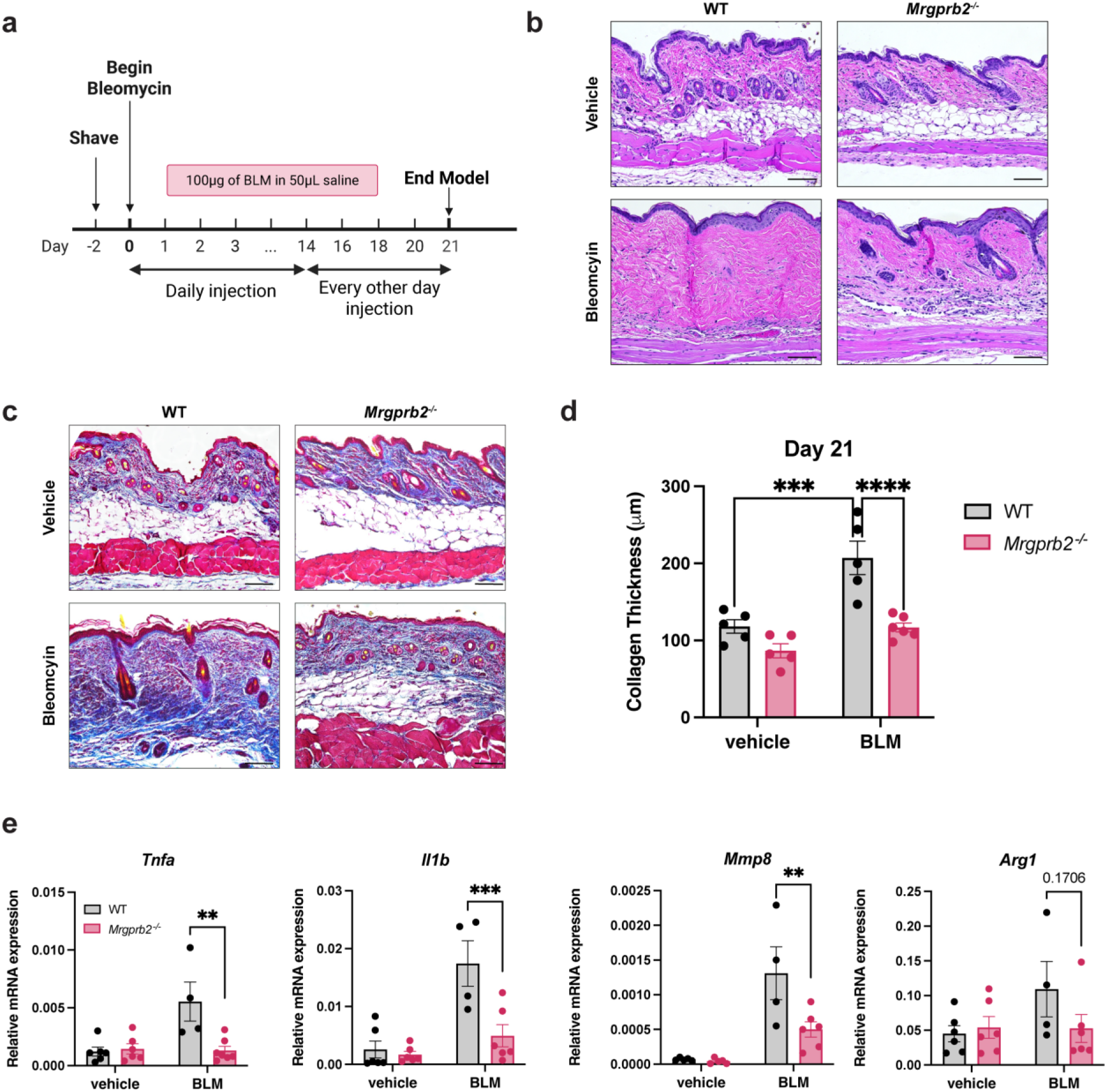
*Mrgprb2*^−/−^ mice are protected from skin fibrosis after BLM treatment. **(a)** Flowchart of BLM injection protocol for inducing skin fibrosis. **(b)** H&E stain of WT and *Mrgprb2*^−/−^ BLM-treated skin at day 21. Scale bar, 100μm. **(c)** Masson’s trichrome stain of WT and *Mrgprb2*^−/−^ BLM-treated skin at day 21. Blue stain indicates collagen. Scale bar, 100μm. **(d)** Quantification of collagen thickness from Masson’s trichrome stain at day 21 (WT veh/BLM *n*=5/5, *Mrgprb2*^−/−^ veh/BLM *n*=5/6). **(e)** Relative mRNA expression normalized to actin in WT/*Mrgprb2*^−/−^ vehicle/BLM skin at day 21 (Tnfa: WT veh/BLM *n*=6/4, *Mrgprb2*^−/−^ veh/BLM *n*=6/7, *Il1b*: WT veh/BLM *n*=6/4, *Mrgprb2*^−/−^ veh/BLM *n*=6/6, *Mmp8*: WT veh/BLM *n*=5/4, *Mrgprb2*^−/−^ veh/BLM *n*=5/6, *Arg1*: WT veh/BLM *n*=5/4, *Mrgprb2*^−/−^ veh/BLM *n*=5/6). Statistical analysis: two-way ANOVA with uncorrected Fisher’s LSD test (D), two-way ANOVA with Sidak’s multiple comparisons test (e). ^**^*P* < 0.01, ^***^*P* < 0.001, ^****^*P* < 0.0001.

As mast cells often mediate innate inflammation, we next assessed proinflammatory markers in the skin using quantitative PCR. *Tnfa* and *Il1b*, genes known to promote inflammatory pathways in keratinocytes^20^ are upregulated in WT-but not *Mrgprb2*^−/−^-treated BLM mice (**Figure 1e**). The matrix metalloproteinase *Mmp8*, associated with SSc and known to remodel the extracellular matrix in the pathogenesis of fibrosis^21,22^, is similarly increased in the skin of WT BLM-treated mice (**Figure 1e**). Given the established role of M2-polarized macrophages in the fibrotic stages of SSc, we also examined Arginase 1 (*Arg1*), a marker of these cells^23,24^. *Arg1* shows a trending increase in WT BLM-treated mice, suggesting a potential activation of fibrotic macrophages in this group (**Figure 1e**). These data indicate that Mrgprb2 plays a role in skin fibrosis through potential inflammatory and matrix remodeling mechanisms.

### Mast cells accumulate in mouse BLM-treated skin and human SSc skin

Given the strong association with Mrgprb2 and collagen deposition, we next hypothesized that mast cells accumulate in the skin of BLM mice in an Mrgprb2-dependent manner. Using the mast cell marker avidin, we observe an increase in mast cell recruitment to the skin of BLM-treated WT compared to *Mrgprb2*^−/−^ mice (**Figure 2a-b**). To rule out the possibility that BLM itself directly stimulates mast cells, we tested whether BLM induces mast cell degranulation via MRGPRX2. BLM does not induce degranulation in either MRGPRX2-expressing or MRGPRX2-knockout cells, confirming that that the effects of the model cannot be explained by direct mast cell activation (**Supplementary Figure 1**).

**Figure 2.**
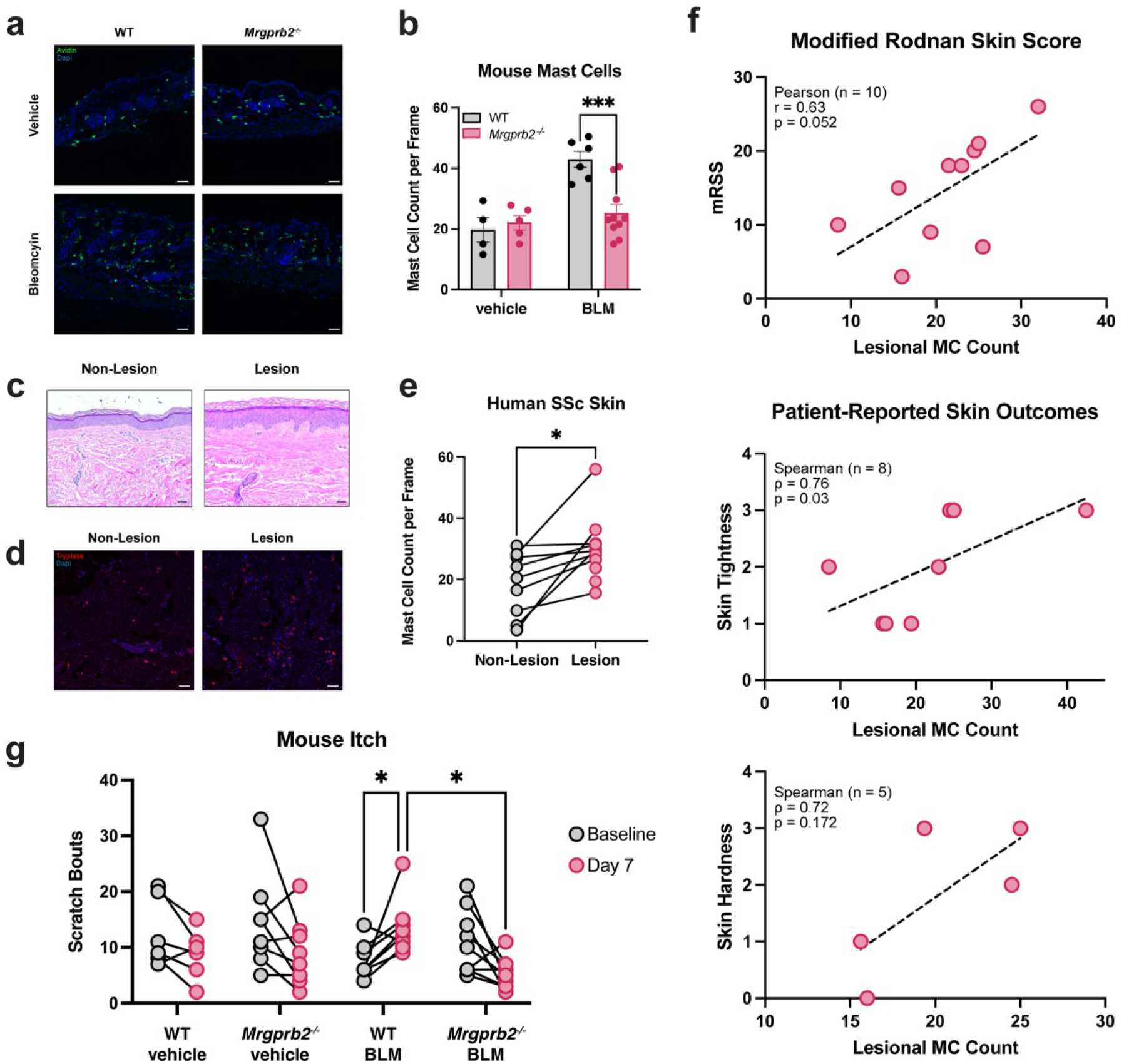
Mast cells accumulate in skin of bleomycin-induced fibrosis in mice and human lesional SSc skin. **(a)** Representative images of avidin stain (green) for mast cells in skin of WT and *Mrgprb2*^−/−^ vehicle- and BLM-treated skin at day 21. Scale bar, 50μm. **(b)** Quantification of mast cell count per frame at day 21 (WT veh/BLM *n*=4/6, *Mrgprb2*^−/−^ veh/BLM *n*=5/10). **(c)** Representative H&E stain of human patient non-lesional and lesional skin. Scale bar, 50μm. **(d)** Representative images of tryptase stain (red) for mast cells in human skin. Scale bar, 50μm. **(e)** Quantification of mast cell count per frame in human patient skin biopsies (non-lesional *n*=9, lesional *n*=11). **(f)** Scatterplots of lesional mast cell count versus (top) mRSS total, (middle) patient-reported skin tightness, and (bottom) patient-reported skin hardness. Each point is one patient, and the dashed line is the least-squares fit. Pearson/spearman correlations and associated p-values are listed. **(g)** Spontaneous scratch bouts during at 30-minute period at baseline and day 7 (WT veh/BLM *n*=6/8, *Mrgprb2*^−/−^ veh/BLM *n*=8/9). Statistical analysis: two-way ANOVA with Sidak’s multiple comparisons test (b), paired Student’s t-test (e), and two-way ANOVA with Tukey’s multiple comparisons test (g). ^*^*P* < 0.05, ^***^*P* < 0.001.

We next wanted to correlate our mast cell findings in the mouse model with human SSc patient skin. We collected lesional and non-lesional skin biopsies from 11 SSc patients and quantified mast cell abundance, as human cutaneous mast cells are known to express MRGPRX2^25^ (**Supplementary Table 1, Figure 2c)**. Across the cohort, lesional skin mast cell counts were consistently higher than paired non-lesional skin counts (**Figure 2d-e**), establishing that mast cells accumulate specifically within affected skin (**Supplementary Figure 2a**). We next investigated whether mast cell burden tracks with the clinical severity of skin disease in SSc. Lesional mast cell count correlated positively with the modified Rodnan skin score (mRSS total; Pearson r = 0.63, p = 0.052, n = 10), a standardized measure of skin thickness used in clinical trials for SSc^26^, where higher scores indicate more severe skin disease. Similarly, lesional mast cell counts correlated more strongly with patient-reported skin tightness (Spearman ρ = 0.76, p = 0.03, n = 8). Patient reported skin hardness followed the same direction but did not reach significance at this sample size (Spearman ρ = 0.72, p = 0.172, n = 5) (**Figure 2f**). Together, these data position lesional skin mast cell burden as a potential tissue correlate of cutaneous disease activity in SSc, though larger cohorts are needed to confirm these associations.

### Mrgprb2 contributes to the development of itch in the BLM model of skin fibrosis

Building on this link between mast cell burden and cutaneous disease activity, we next asked whether mast cells also contribute to the pathogenesis of pruritus in SSc. Since itch is often an early symptom of disease^4^, we quantified spontaneous scratching bouts of WT and *Mrgprb2*^−/−^ mice during the early stage of BLM treatment, within the first seven days. By day 7, spontaneous scratching is increased in BLM-treated WT mice (**Figure 2g, Supplementary Figure 2a**). BLM-treated *Mrgprb2*^−/−^ mice do not show this same increase and have significantly fewer scratching bouts on day 7 compared to WT mice. These data suggest that early pruritus in this model may be associated with mast cell infiltration and activity, nominating mast cell burden as a candidate link to cutaneous disease severity in SSc and warranting further investigation into its relationship with patient-reported itch.

## Discussion

The role of mast cells in SSc has been largely understudied, in part due to unclear pathways of activation. Here, we show that mast cells, via Mrgprb2, contribute to skin fibrosis in a BLM mouse model. *Mrgprb2*^−/−^ mice show reduced mast cell accumulation and markedly less collagen deposition compared to WT mice, alongside lower induction of inflammatory and fibrotic factors. Consistent with this, human SSc lesional skin harbors significantly more mast cells than non-lesional skin, and lesional mast cell counts trended with greater global skin severity as reflected in mRSS and patient-reported skin tightness and hardness.

In addition to contributing to the classic fibrotic features of SSc, Mrgprb2 also contributes to early pruritus in the BLM mouse model, the first report of pruritus in this model. These distinct kinetics of early-stage pruritus versus late-stage fibrosis may reflect a transition from acute to chronic mast cell activation phenotypes. Early pruritus likely arises from initial mast cell degranulation and pruritogen release, while persistent activation in later disease instead drives immune recruitment and matrix remodeling. Some data suggest that chronic cytokine exposure can desensitize and downregulate mast cell receptor expression and activity, altering mast cell degranulation in chronic inflammatory conditions^27^. This may explain why early mast cell degranulation-induced pruritus often precedes chronic, mast cell-enhanced fibrosis in SSc and raises the possibility that mast cell-stabilizing interventions could blunt itch in early disease while requiring distinct, chronic inhibition for fibrosis-directed therapy. Together, these data suggest a role for Mrgprb2 in the exacerbation of fibrosis and pruritus in SSc and point to a broader, previously underappreciated role for mast cells in autoimmune connective tissue disease.

## Limitations

This study has key limitations that warrant further investigation into the relationship between mast cell burden and SSc cutaneous disease. While we correlate mast cell counts with progression of skin fibrosis and pruritus in mouse models, these experiments rely on bleomycin-induced fibrosis rather than the environmental and genetic drivers likely responsible for human SSc. Further, the patient cohort’s limited sample size restricts our ability to stratify by disease subtype and duration, and correlations between mast cell counts and disease features should be interpreted with this constraint in mind. Further studies with larger, more diverse patient cohorts will broaden our understanding of the role of mast cells in SSc disease burden.

## ABBREVIATIONS

Mrgprb2: (Mas-related G protein-coupled receptor b2)
MRGPRX2: (Mas-related G protein-coupled receptor X2)
SSc: (systemic sclerosis)
BLM: (bleomycin)
Tnfa: (tumor necrosis factor alpha)
Il1b: (interleukin 1 beta)
Arg1: (arginase 1)
mRSS: (modified Rodnan skin score)

## Ethics Statement

This study was performed after approval by the Johns Hopkins Institutional Animal Care and Use Committee and Institutional Review Board for animal experiments and patient sample analyses, respectively.

## Data Availability

The authors confirm that the data presented within this study are available within the figures, tables, and supplementary data files.

## Acknowledgements

Illustrations/diagrams were created using BioRender. We thank Dean D. Metcalfe from the National Institute of Allergy and Infectious Diseases for providing us with LAD2 mast cells. The work was supported by the Howard Hughes Medical Institute to X.D. X.D is also supported by R37NS054791. JK is supported by the Dermatology Foundation Medical Dermatology Career Development Award. AAS is supported by K24AR08217 and The Sara and Alex Othon Research Fund.

## Author Contributions

Conceptualization: RK, JK, XD; Investigation: RK, DC, CK, HO, SP, AS, GL, AS; Formal Analysis: RK; Funding acquisition: XD; Writing - Original Draft Preparation: RK; Writing - Review and Editing: RK, CK, JK, XD.

## Declaration of Generative AI Use

The authors did not use AI/LLM in any part of the research process or manuscript preparation.

## Methods

### Bleomycin Model

Mice were shaved one day prior to induction of the model. Mice were anesthetized and injected with 100μg of BLM (in 50μL of saline) or vehicle (50μL of saline) subcutaneously in the nape of the neck every day for 14 days, followed by every other day injections until day 21.

### Itch Behavior

Mice were shaved and habituated two days before itch recording by placement in the itch chamber for 1 hour each day. Baseline recordings were taken before induction of BLM injections at day 0. Recordings were taken for 30 minutes every day in the morning after a 15-minute habituation period. Injections were done 4 hours after recordings such that spontaneous itch was measured rather than acute-induced itch. Each 30-minute recording was counted for scratching bouts.

### RNA isolation and quantification

Skin was carefully harvested from WT and *Mrgprb2*^−/−^ adult mice. Total RNA was extracted using the RNeasy® Plus Micro Kit (Qiagen), according to the manufacturer’s instructions. RNA was quantified using NanoDrop. Reverse transcription was performed using the iScript cDNA synthesis kit and following the manufacturer’s suggestions. Finally, qPCR was performed using TaqMan™ Fast Advanced Master Mix (Applied Biosystems) and run on a StepOnePlus Real-Time PCR System (Applied Biosciences) and analyzed by StepOne Software v2.2.2.

### Immunofluorescence

#### Mouse and Human Skin

Adult mice were sacrificed using CO_2_. Skin biopsies were collected and placed in 4% paraformaldehyde (PFA) at 4°C overnight, followed by embedding in paraffin. After sectioning, samples were deparaffinized. Sections were washed with PBS, then placed in blocking solution (10% normal goat serum in PBS, 0.2% Triton-X) for 1 h at room temperature. Sections were then washed with PBS (3 washes, 5 min each) and incubated in avidin (1:500) in blocking solution at room temperature for 2h. Slides were then washed with PBS (5 washes, 5 min each), mounted with Fluoromount-G with DAPI (Invitrogen), and imaged using the Zeiss LSM700 Confocal microscope.

For human skin biopsies, samples were placed in formalin and embedded in paraffin for sectioning. After sectioning with a 5μm thickness, samples were deparaffinized using xylene and ethanol washes, followed by antigen retrieval in 1X RNAScope Target Retrieval reagent (Advanced Cell Diagnostics) for 15 minutes in a steamer held at 99°C. Slides were then dehydrated in 100% ethanol, followed by blocking and staining as above. Sections were incubated in antibody overnight in blocking solution at 4°C as follows: anti-tryptase (1:500). Sections were then washed with PBS (3 washes, 5 min each) and incubated in secondary antibody in blocking solution at room temperature for 2h. All secondary antibodies (Life Technologies) were diluted 1:500. Slides were then washed with PBS (5 washes, 5 min each), mounted with Fluoromount-G with DAPI (Invitrogen), and imaged using the Zeiss LSM700 Confocal microscope. Representative mast cell counts were determined by individuals blinded to phenotype and counts were reported as an average across three sections per sample.

### H&E and Masson’s Trichrome Stain

Sections were submitted to the Johns Hopkins Reference Histology Core for H&E staining. For Masson’s Trichrome stain, a commercially available kit was used according to manufacturer’s instructions (Abcam). Slides were mounted with synthetic resin and imaged using the Nikon Eclipse TE200. Images were then blinded and collagen thickness was measured on the microscope as the average thickness across three sections per sample.

### Beta Hexosaminidase Degranulation Assay

WT and MRGPRX2^-/-^ LAD2 mast cells were seeded at 5 x 10^5^ cells/mL per well in a 96-well round bottom plate. 50μL of 2X treatment was then added to the cells, to obtain a final 1X concentration in the well. Substance P was treated using a 100nM final concentration, and bleomycin was treated at a final concentration of 50μM. Cells were treated for 30 minutes then spun down at 300g for 5 min. 50μL of supernatant and 50μL of cell lysate (0.1% Triton-X in PBS) were then added to 50μL of p-nitrophenyl N-acetyl-β-D-glucosaminide (Sigma-Aldrich) in 0.1 M sodium citrate buffer (pH 4.5) for 90 minutes at 37°C. After incubation, the reaction was stopped by adding 50μL of 0.4M pH 10.7 glycine buffer to each well. The plate was immediately read at absorbance 405nm with 570nm as reference. The ratio of beta hexosaminidase released was calculated by the following formula:

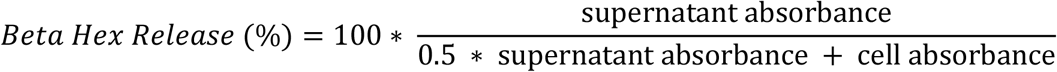

### Patient Cohort Review and Statistical Analysis

In this cross-sectional study, adults with systemic sclerosis (SSc) evaluated by the departments of dermatology and rheumatology at the Johns Hopkins Hospital, were recruited under approval by the Johns Hopkins Institutional Review Board. Skin samples were obtained as formalin-fixed, paraffin-embedded (FFPE) blocks from two biorepositories: the Johns Hopkins Scleroderma Center Cutaneous Tissue Repository (NA00070546) from the department of Rheumatology, and the *Cutaneous Autoimmune Connective Tissue Disease Database and Biorepository (IRB00339015) from the department of Dermatology*. All patients were adults greater than 18 years of age, who met the 2013 ACR/EULAR classification criteria for SSc. Patients were excluded for absence of a skin biopsy available for mast cell quantification. Where both lesional and non-lesional biopsies were available, samples were analyzed as matched pairs; patients with a lesional biopsy only were retained for lesional-count analyses.

We collected sociodemographic characteristics (age at biopsy, sex, race, ethnicity), disease characteristics (SSc subtype, myositis-specific and SSc-associated autoantibodies, smoking history), and cutaneous and systemic disease measures from chart review, including the modified Rodnan skin score (mRSS), pulmonary function, and patient-reported skin symptoms (tightness, hardness, pain, redness, and stiffness). Staining of skin samples was carried out as mentioned above. Patient-reported skin symptoms were recorded on ordinal descriptor scales (e.g., “not at all” through “severe”) and recoded to ordered numeric scores (0–4) for analysis; values coded as missing were excluded from the respective analyses.

Statistical analyses were performed in Stata 18.0 (StataCorp LLC). Lesional versus non-lesional mast-cell counts were compared in the 9 paired patients. The lesional dermal mast-cell count was the primary predictor of interest, tested against each disease feature as the outcome: continuous outcomes (mRSS) were assessed with Pearson correlation and linear regression, ordinal skin-symptom scores with Spearman rank correlation, and binary features with logistic regression. As the sample was too small to support a full multivariable model, adjustment was with each of the following singly covariates – age at biopsy, sex, race, ethnicity, SSc subtype, and smoking history, using multivariable linear regression for continuous outcomes and partial (rank-based) correlation for ordinal outcomes.

### Quantification and Statistical Analysis

Statistical analyses other than the patient chart review were performed using GraphPad Prism v.10.0.2. Statistical comparisons were conducted by two-tailed, unpaired Student’s t-test, or two-way ANOVA with Sidak’s multiple comparison test or Tukey’s multiple comparison test, unless otherwise noted. For all bar charts, bars depict the mean. All error bars represent SEM and n represents the number of mice analyzed. Specific details of each experiment, including statistical test used and exact n value, can be found in the figure legends.

**Supplementary Figure 1.**
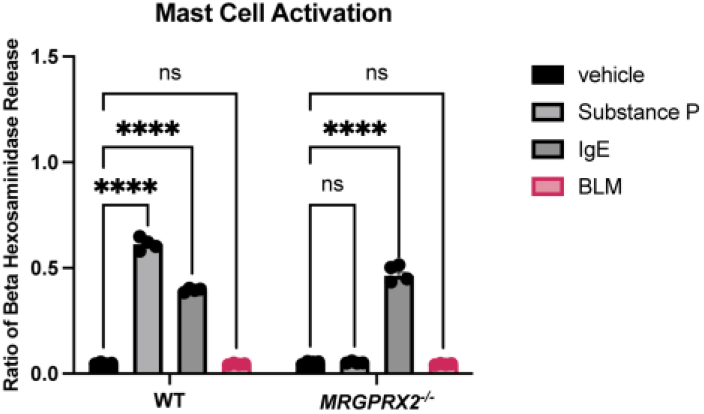
BLM does not activate MRGPRX2. WT and *MRGPRX2*^-/-^ LAD2 human mast cell beta-hexosaminidase release after BLM treatment, using SP and IgE as positive controls (*n=*4 biological replicates for all conditions). Statistical analysis: two-way ANOVA with Sidak’s multiple comparisons test (A-B). ^****^*P* < 0.0001.

**Supplementary Figure 2.**
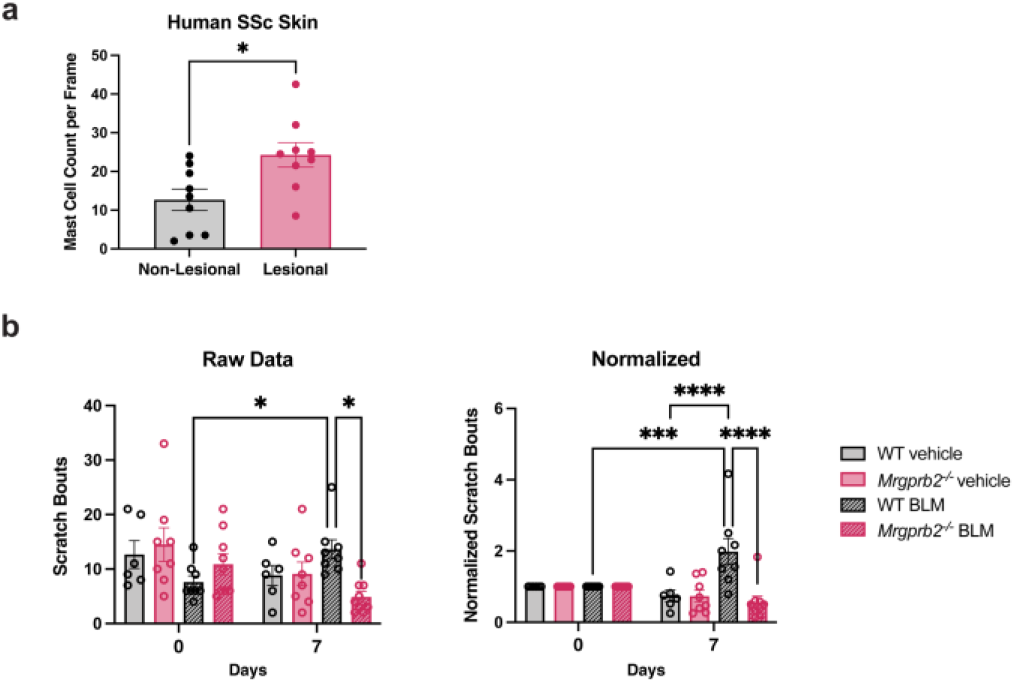
Mrgprb2 contributes to itch in the bleomycin model of skin fibrosis. **(a)** Same data as seen in Figure 2e, plotted as pooled non-lesional and lesional samples with no pairing, and with bar graphs to outline means and SEM. **(b)** Same data as seen in Figure 2g, plotted in bar graph format to outline means and SEM. (left) Raw data and (right) data normalized to baseline itch counts for each mouse. Statistical analysis: two-sided Student’s t-test (a), two-way ANOVA with Tukey’s multiple comparisons test (b). ^*^*P* < 0.05, ^***^*P* < 0.001, ^****^*P* < 0.0001.

**Supplementary Table 1.** Demographic, Clinical, and Serologic Features of Patients with SSc (n = 11)

| **Characteristic** | **n (%)** |
| --- | --- |
| ***Demographics*** |  |
| Age at biopsy, mean (SD), y | 51.3 (10.8) |
| Duration of disease at biopsy, mean (SD), y | 1.8 (2.1) |
| Female sex, n (%) | 9 (81.8) |
| Race |  |
| *White race, n (%)* | 7 (63.6) |
| *Black race, n (%)* | 2 (18.2) |
| Ethnicity, n (%) |  |
| *Hispanic, n (%)* | 2 (18.2) |
| Smoking History |  |
| *Current, n (%)* | 2 (18.2) |
| *Former, n (%)* | 3 (27.3) |
| Family history of autoimmune disease, n (%) | 4 (36.4) |
| ***Comorbidities*** |  |
| Hypertension, n (%) <sup>a</sup> | 7 (63.6) |
| Dyslipidemia, n (%) <sup>b</sup> | 3 (27.3) |
| COPD, n (%) <sup>a</sup> | 2 (18.2) |
| ***SSc Disease Features (at time of biopsy)*** |  |
| Diffuse cutaneous subtype, n (%) | 9 (81.8) |
| Limited cutaneous subtype, n (%) | 1 (9.1) |
| Raynaud phenomenon, n (%) | 7 (63.6) |
| Modified Rodnan skin score, mean (SD) <sup>a</sup> | 14.7 (7.2) |
| Scleroderma renal crisis, n (%) | 1 (9.1) |
| ***Vascular and Cutaneous Involvement (at time of biopsy)*** |  |
| Digital ulcers, n (%) | 1 (9.1) |
| Digital pitting, n (%) | 2 (18.2) |
| Telangiectasias, n (%) | 5 (45.5) |
| Calcinosis, n (%) | 1 (9.1) |
| ***Systemic Involvement*** |  |
| Dyspnea, n (%) <sup>a</sup> | 7 (63.6) |
| Arthralgia, n (%) | 5 (45.5) |
| ***Autoantibodies*** |  |
| ANA, n (%) | 9 (81.8) |
| Anti-centromere, n (%) | 1 (9.1) |
| Anti- Scl-70 (topoisomerase I), n (%) | 2 (18.2) |
| Anti-RNA polymerase III, n (%) | 4 (36.4) |
| ***Pulmonary Function (closest to time of biopsy)*** |  |
| FVC, % predicted, mean (SD) <sup>a</sup> | 87.7 (16.8) |
| DLCO, % predicted, mean (SD) <sup>b</sup> | 67.1 (17.2) |
| Pulmonary hypertension, n (%) | 1 (9.1) |
| ***Treatment*** |  |
| Any immunomodulatory therapy, n (%) | 9 (81.8) |
| *Hydroxychloroquine, n (%)* | 2 (18.8) |
| *IVIg, n (%)* | 2 (18.8) |
| *Methotrexate, n (%)* | 1 (9.1) |
| *Mycophenolate Mofetil, n (%)* | 6 (54.5) |
| *Prednisone, n (%)* | 1 (9.1) |
| *Rituximab, n (%)* | 1 (9.1) |

Abbreviations: COPD, chronic obstructive pulmonary disease; ANA, antinuclear antibody; FVC, forced vital capacity; DLCO, diffusing capacity of the lungs for carbon monoxide; IVIG, intravenous immunoglobulin. Duration of disease at biopsy is from the time of disease diagnosis per chart review. Treatments indicates use of these drugs at any point for SSc up until time of biopsy.
<sup>a</sup> Data missing for 1 patient.
<sup>b</sup> Data missing for 2 patients.
Percentages are calculated among patients with non-missing data. Continuous variables are shown as mean (SD); categorical variables as n (%).

**Supplementary Table 2.**
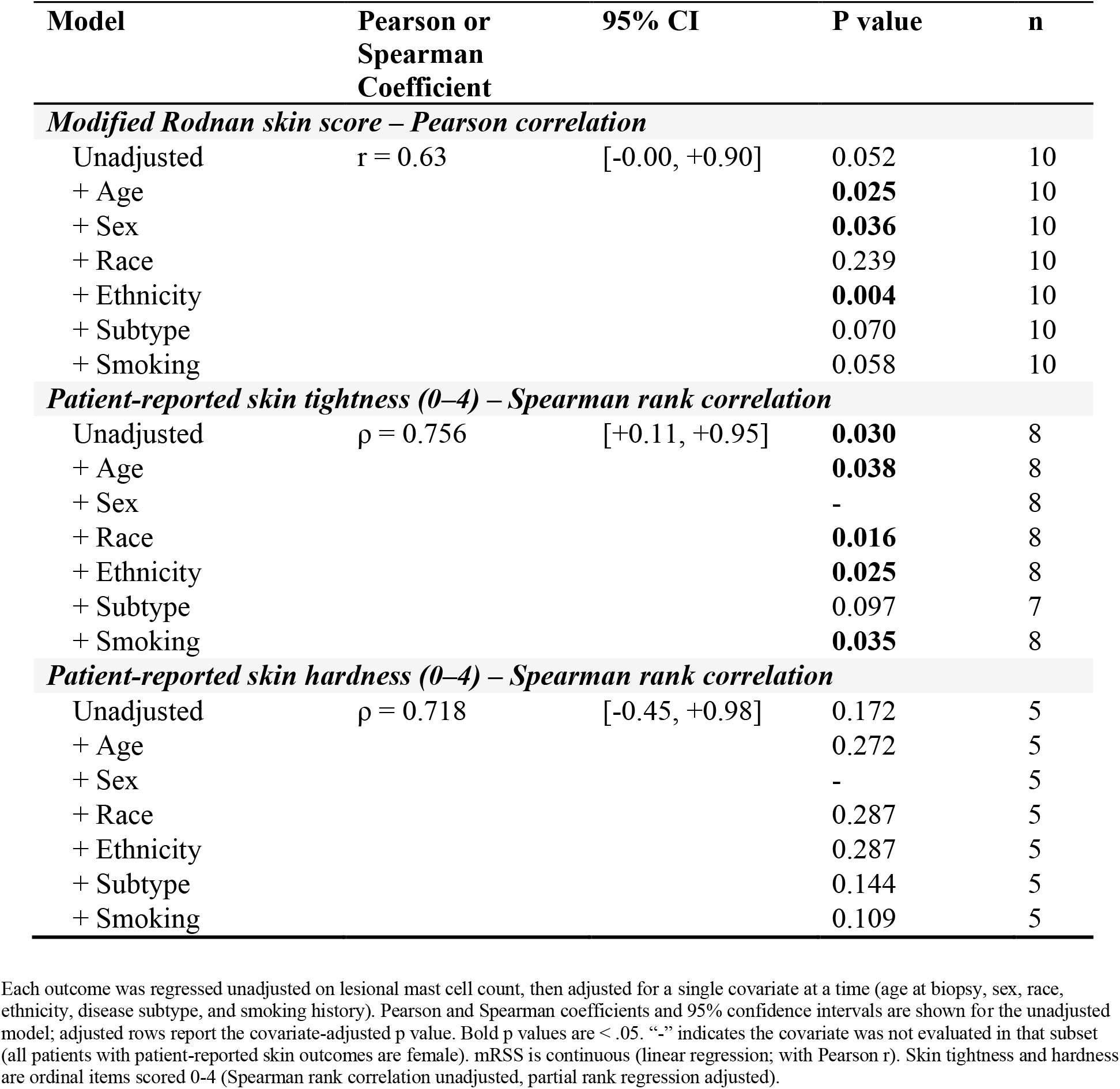
Regression of Skin Severity Outcomes and Lesional Mast Cell Counts (n = 11)

| Model | Pearson or Spearman Coefficient | 95% CI | P value | n |
| --- | --- | --- | --- | --- |
| ***Modified Rodnan skin score – Pearson correlation*** |  |  |  |  |
| Unadjusted | r = 0.63 | [-0.00, +0.90] | 0.052 | 10 |
| + Age |  |  | **0.025** | 10 |
| + Sex |  |  | **0.036** | 10 |
| + Race |  |  | 0.239 | 10 |
| + Ethnicity |  |  | **0.004** | 10 |
| + Subtype |  |  | 0.070 | 10 |
| + Smoking |  |  | 0.058 | 10 |
| ***Patient-reported skin tightness (0–4) – Spearman rank correlation*** |  |  |  |  |
| Unadjusted | $\rho = 0.756$ | [+0.11, +0.95] | **0.030** | 8 |
| + Age |  |  | **0.038** | 8 |
| + Sex |  |  | - | 8 |
| + Race |  |  | **0.016** | 8 |
| + Ethnicity |  |  | **0.025** | 8 |
| + Subtype |  |  | 0.097 | 7 |
| + Smoking |  |  | **0.035** | 8 |
| ***Patient-reported skin hardness (0–4) – Spearman rank correlation*** |  |  |  |  |
| Unadjusted | $\rho = 0.718$ | [-0.45, +0.98] | 0.172 | 5 |
| + Age |  |  | 0.272 | 5 |
| + Sex |  |  | - | 5 |
| + Race |  |  | 0.287 | 5 |
| + Ethnicity |  |  | 0.287 | 5 |
| + Subtype |  |  | 0.144 | 5 |
| + Smoking |  |  | 0.109 | 5 |

Each outcome was regressed unadjusted on lesional mast cell count, then adjusted for a single covariate at a time (age at biopsy, sex, race, ethnicity, disease subtype, and smoking history). Pearson and Spearman coefficients and 95% confidence intervals are shown for the unadjusted model; adjusted rows report the covariate-adjusted p value. Bold p values are < .05. “-” indicates the covariate was not evaluated in that subset (all patients with patient-reported skin outcomes are female). mRSS is continuous (linear regression; with Pearson r). Skin tightness and hardness are ordinal items scored 0–4 (Spearman rank correlation unadjusted, partial rank regression adjusted).

